# Allostery in STAT3 Variant D170A is Mediated by a Rigid Core

**DOI:** 10.1101/2022.06.15.495314

**Authors:** Tingting Zhao, Nischal Karki, Brian Zoltowski, Devin A. Matthews

## Abstract

Signal Transducer and Activator of Transcription 3 (STAT3) plays a crucial role in cancer development and thus is a viable target for cancer treatment. STAT3 functions as a dimer mediated by phosphorylation of the SRC-homology 2 (SH2) domain, a key target for therapeutic drugs. While great efforts have been employed towards the development of compounds that directly target the SH2 domain, no compound has yet been approved by the FDA due to a lack of specificity and pharmacologic efficacy. Studies have shown that allosteric regulation of SH2 via the coiled-coil domain (CCD) is an alternative drug design strategy. Several CCD effectors have been shown to modulate SH2 binding and affinity, and at the time of writing at least one drug candidate has entered phase I clinical trials. However, the mechanism for SH2 regulation via CCD is poorly understood. Here, we investigate structural and dynamic features of STAT3 and compare the wild type to the reduced function variant D170A in order to delineate mechanistic differences and propose allosteric pathways. Molecular dynamics simulations were employed to explore conformational space of STAT3 and the variant, followed by structural, conformation, and dynamic analysis. The trajectories explored show distinctive conformational changes in the SH2 domain for the D170A variant, indicating long range allosteric effects. Multiple analyses provide evidence for long range communication pathways between the two STAT3 domains, which seem to be mediated by a rigid core which connects the CCD and SH2 domains via the linker domain (LD) and transmits conformational changes through a network of short-range interactions. The proposed allosteric mechanism provides new insight into the understanding of intramolecular signaling in STAT3 and potential pharmaceutical control of STAT3 specificity and activity.

**Author Summary:** In all living organisms, the proliferation and survival of cells are regulated by various proteins. Signal Transducers and Activators of Transcription 3(STAT3) protein is one of the important proteins. However, the abnormal regulation of these proteins will lead to cancer cell. The constitutive activation of STAT3 has been linked to several types of solid tumors, leukemia, and lymphomas. Consequently, STAT3 proteins have been a key target for cancer therapy. SH2(SRC-homology 2) domain is the key interaction site, great efforts have been attributed to target SH2 domain, which specificity has been a major challenge in drug discovery. Research showing regulation of SH2 domain via CCD has opened a new path for drug discovery, however is challenged by poor understanding of the allosteric mechanism. Here, we show that CCD regulates SH2 conformation via a rigid backbone. The perturbations in CCD is transmitted through α-helix to the rigid core that concert the movement of CCD and LD (Link domain), leading to structural changes in the SH2 domain. The present findings provide allosteric mechanism with atomistic details underlying the regulation of CCD to SH2 domain in STAT3 protein. Which allows informed drug design targeting CCD for desired downstream effect on SH2 domain and the overall STAT3 function.

## Introduction

Proteins within the Signal Transducers and Activators of Transcription (STAT) family function as both signal transducers in the cytoplasm and transcription factors upon nuclear translocation. All members of STAT family consists of six domains (Figure 1A): aminoterminal domain (), coiled-coil domain (CCD), DNA-binding domain (DBD), linker domain (LD), SRC-homology 2 domain (SH2), and transactivation domain (TAD) which is also named the C terminal domain (Lim and Cao ^1^). STAT proteins are regulated by Janus Kinases (JAKs) where they play a crucial role in immune response, cell division, and apoptosis, as a gene expression regulatory arm of JAK-STAT signaling pathway (Aaronson and Horvath ^2^). However, each member is activated via different types of cytokines and have unique function in the pathway(Avalle et al. ^3^, Rani and Murphy ^4^, Stritesky and Kaplan ^5^).

**Figure 1:**
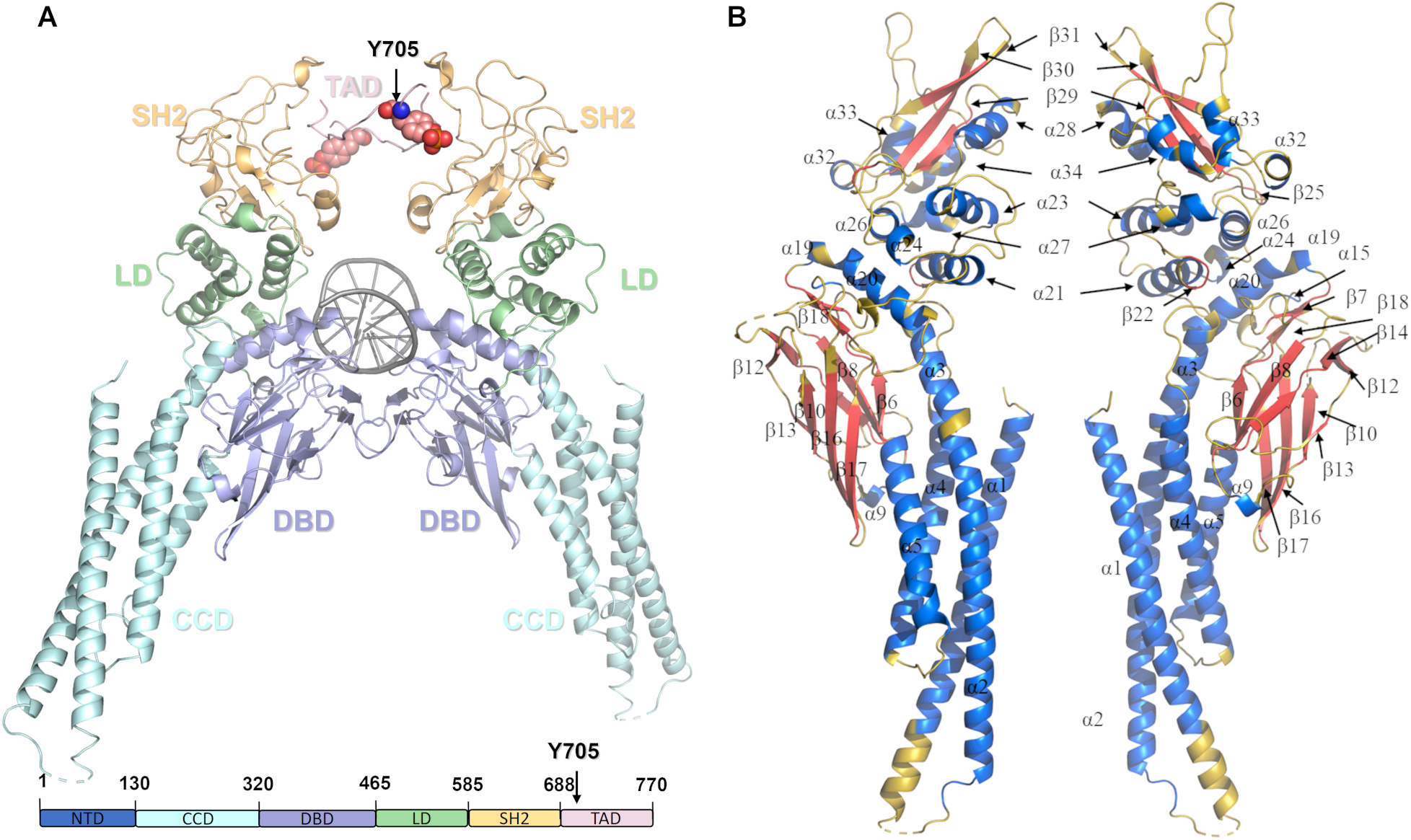
STAT3 structure. **(A)** STAT3 Domain structure (1BG1 as template, NTD not shown, Y705 is shown as spheres). **(B)** Secondary structures are labeled according to the UniProt database (Table S1) (Bateman et al. ^6^): α helices are colored blue, β sheets are colored red, and unstructured regions (loops) are colored yellow (transverse view).

Constitutive activation of STAT3 has been shown to play a crucial role in cancer progression(Sgrignani et al. ^7^, Yu and Jove ^8^). STAT3 binds, via the SH2 domain, to cell-surface receptors upon activation and recruitment of receptor-associated kinases. Upon binding, the recruited kinases activate STAT3 through phosphorylation within the TAD (at Y705), followed by dissociation from the receptor to form homodimers through reciprocal interactions between the SH2 domain and the phosphotyrosine (pY705) residue. These activated homodimers are then translocated to the nucleus where the DBD binds to target genes, and TAD activates the expression of proteins crucial for cell growth and survival. In normal cells, the signaling pathway is well-regulated. However, the abnormal activation of this signaling pathway promotes the development of cancer: misregulation of STAT3 in cancer cells promotes pro-oncogenic inflammation and suppresses anti-tumor immunity (Yu et al. ^9^).

The direct therapeutic inhibition of STAT3 is highly desirable but remains challenging as evident from the lack of FDA–approved drugs. Specifically, significant amounts of effort has been employed to develop molecules targeting the SH2 domain (Johnson et al. ^10^, Bai et al. ^11^, Arshad et al. ^12^). The SH2 domain is a structurally conserved protein domain, which appears in many intracellular signal transducing proteins, offering a binding site for phosphorylated tyrosine residues. The domain contains two regions with specialized functions: the pY pocket, into which the phosphotyrosine of the target inserts, is the binding region, while residues of the pY+3 pocket interact with the three C-terminal residues of the phosphotyrosine in the target, forming a specificity–determining region (Bradshaw et al. ^13^, Haan et al. ^14^). The inhibitors targeting the SH2 domain: phosphotyrosine motifs (pY-peptide) or phosphotyrosine-based peptidomimetic inhibitors which mimic the pTyr-Xaa-Yaa-Gln motif, have been previously investigated, as well as the associated binding mode (Dhanik et al. ^15^, Mandal et al. ^16^, McMurray ^17^). In the pY pocket, R609 is the principal binding partner, along with K591, S636 and S611 which directly interact with pY705. The relative conformation and position of these residues will have a direct effect on STAT3 binding activity. In the pY+3 pocket, V637 in β31 controls accessibility to this pocket, while Y657, Q644, Y640, and E638 facilitate the hydrogen bond interaction with its target, as well as I659, W623 and F621 assist in binding of target peptide by forming hydrophobic environment(McMurray ^17^). However most of these compounds have yet to be explored in clinical studies or further development of these compounds was limited due to concerns with their relative lack of potency and selectivity(Gelain et al. ^18^).

Several studies (Minus et al. ^20^, Huang et al. ^21^, Sala et al. ^22^) have determined that effector (small molecule and polypeptide) binding to CCD interferes with SH2 domain binding or precludes STAT3 nuclear translocation, which suggests potential targets for further drug design. The coiled-coil domain is considered to be essential for STAT3 recruitment to the receptor. Systematic deletion analysis of the N-domain and α helices of CCD, as well as mutagenesis of conserved residues (D170A) in the CCD of STAT3 showed the diminishment of both pY-peptide binding and tyrosine phosphorylation (Zhang et al. ^19^). Furthermore, the small molecule MM-206 was identified as an inhibitor of STAT3 phosphorylation, and Minus et al. surprisingly found the binding site was at α1 of CCD (around F174). In addition, the compound K116, found to bind to CCD by AlloFinder, was shown to be able to inhibit receptor binding, validated by mutagenesis and functional experiments (Huang et al. ^21^). Recently, a small polypeptide MS3-6 was found to bind to CCD and to diminish DNA binding and nuclear translocation. Interaction with the IL-22 receptor was disrupted, but binding to pY-peptide was maintained. These observations are summarized in Table 1. The discovery of several and diverse inhibitory agents, which bind to the CCD rather than SH2 is a fascinating development. However, rational design of allosteric effectors requires a more detailed, mechanistic knowledge of how CCD binding affects SH2 structure and activity. In this work, we study the allosteric mechanism of D170A mutation on the inhibition of STAT3 activity as predicted by the dynamic structures of the SH2 domain known to be essential for pY-peptide binding and ensuing Y705 phosphorylation. Specifically, we investigate the structural properties within the SH2 domain upon CCD mutation over a number of molecular dynamics simulations, as well as the dynamical correlations between SH2 and CCD and the associated networks of allosteric residues and interactions within various structural motifs.

**Table 1:** Summary of STAT3 effectors targeting CCD and their effect on pY-peptide and DNA binding activity. a) Zhang et al. ^19^, b)Minus et al. ^20^, c)Huang et al. ^21^, d)Sala et al. ^22^

| Alteration/Effector | pY-Peptide Binding Activity |  | DNA Binding Activity |
| --- | --- | --- | --- |
|  | Sequence / Activity (SPR-based assay) | Phosphorylation Agent / Activity (western blot) |  |
| D170A mutation<sup>a)</sup> | VVHSG(pY)RHQVPS / Inhibited | EGF / Inhibited | Not studied |
| MM-206<sup>b)</sup> | LPVPE(pY)INQSVP / Inhibited | IL-6 / Inhibited | Inhibited |
| K116<sup>c)</sup> | Ac-pYLPQTV-NH<sub>2</sub> / Inhibited | IL-6 induced / Inhibited | Not studied |
| MS3-6<sup>d)</sup> | GMPKS(pY)LPQTVR / Not inhibited | IL-22 / Inhibited<br>IL-6 / Not inhibited | Diminished |

Q644, Y640, and E638 facilitate the hydrogen bond interaction with its target, as well as I659, W623 and F621 assist in binding of target peptide by forming hydrophobic environment (McMurray<sup>17</sup>). However most of these compounds have yet to be explored in clinical studies or further development of these compounds was limited due to concerns with their relative lack of potency and selectivity (Gelain et al.<sup>18</sup>).

## Results

### D170A mutation induces large structural but minor dynamical changes in STAT3

The MD trajectories for both wild type and D170A variants were analyzed using RMSD. To reconcile the change in system from NPT to NVT, RMSD was computed by using the first frame of the NVT system as the reference. Both the structures reached an equilibrium after 100 ns, where the average RMSD stabilized around 4 with respect to the starting structure (Figure S1). The RMSD of each domain was also calculated and shown in (Figure S1), the large mobility indicated by RMSD was mainly contributed by the SH2 domain. An RMSD cross-correlation analysis between different domains shows a distinct increase in correlation between the CCD and SH2 domains in the D170A variant compared to the wild-type (Figure 2 A, 2 B). This correlation is enhanced at the expense of correlation of different domains to the linker domain, with only CCD retaining its correlations with linker in D170A variant.

**Figure 2:**
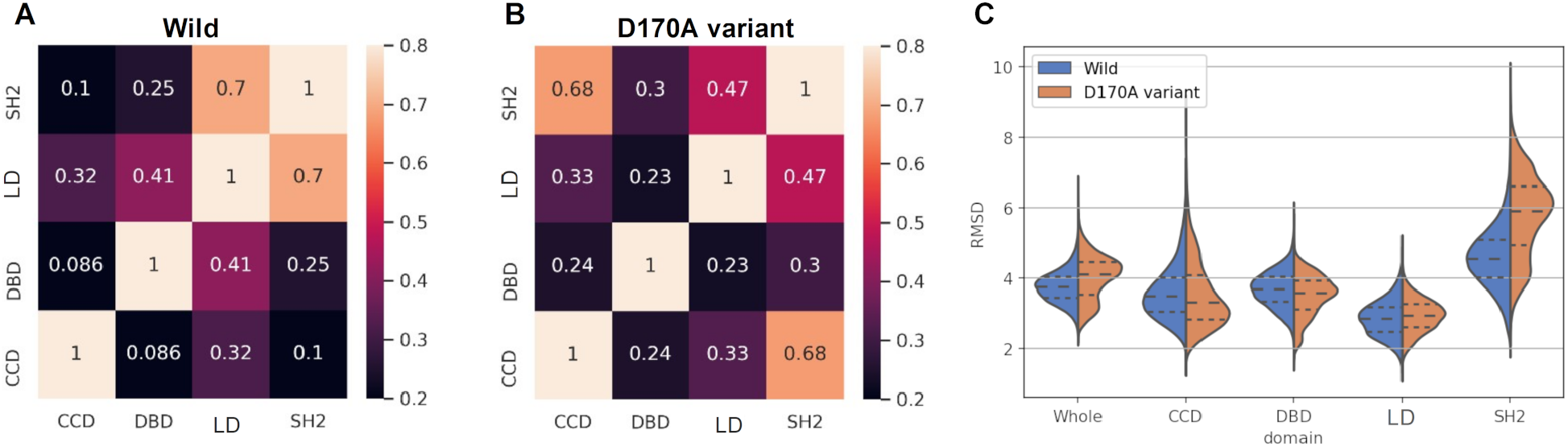
Root Mean Square Deviation (RMSD) analysis. **(A, B)** Cross-correlation (Pearson correlation) of RMSD values of each domain for wild type and D170A variant. Here RMSD values were calculated with the first frame of reference since the correlation of dynamic changes of each domain is of interested. While worth being noted, the correlation value does not suggest the functional correlation between domains, since RMSD is an overall measurement of conformational changes with regarding to the reference structure, distinctive conformations may have same RMSD value. **(C)** Violin plot of RMSD values for the whole protein (core full length protein, not including NTD) and each domain. Here crystal structure was using as reference, since the conformational changes difference between wild type and D170A was of interested.

Using the crystal structure (PDB ID: 6TLC) as the reference, RMSD values of wild type and the D170A variant were computed and shown in a violin plot (Figure 2 C). The distribution of RMSD values demonstrates that the D170A variant has multiple quasi-stable conformations seen as multiple peaks in the core full length protein violin plot, whereas the wild type consists of a single pronounced peak. Investigation of each domain shows similar trends, with SH2 having the largest difference between wild type and mutant due to significantly higher RMSD values in D170A, and with LD showing the least change upon mutation. Overall, the deviation of the SH2 domain from the crystal structure is significantly increased and several quasi-stable conformations appear, while the CCD domain becomes somewhat more rigid. However, the CCD domain also exhibits a minority configuration with increased deviation from the crystal structure. This minority configuration corresponds to a kinked configuration of the α1 helix, as is described in our clustering analysis (Section 2.3).

RMSF analysis shows that the difference in conformational dynamics between the wild type and the D170A variant is statistically insignificant in our sample set, even though the overall structural conformations of the two variants are quite different. Qualitative observation of the mean RMSF values shows only slight differences in each of the domains.

Specifically, small increases in flexibility are seen in the α1–α2 and α4–α5 loops in CCD, as well as the β11, β11–β12, β13–β14, and β14–α15 loops in DBD. Inversely, flexibility in the D170A mutant is slightly decreased in the β22, α23–α24 loop, α24–β25 loop, and β25 in LD, and the β30–β31 and α33–34 loops in SH2 domain (Figure 3). From these qualitative observations, it can be inferred that the increased flexibility in CCD due to loss of a negatively charged residue D170, leads to an allosteric increase in flexibility of the DBD and a decrease in flexibility of the LD and SH2 domains. Even though the changes in overall flexibility are small, the D170A variant experiences significant changes in the number and geometry of the available quasi-stable conformations within each domain as well as the whole protein. The source of these conformational changes is explored in the following sections.

**Figure 3:**
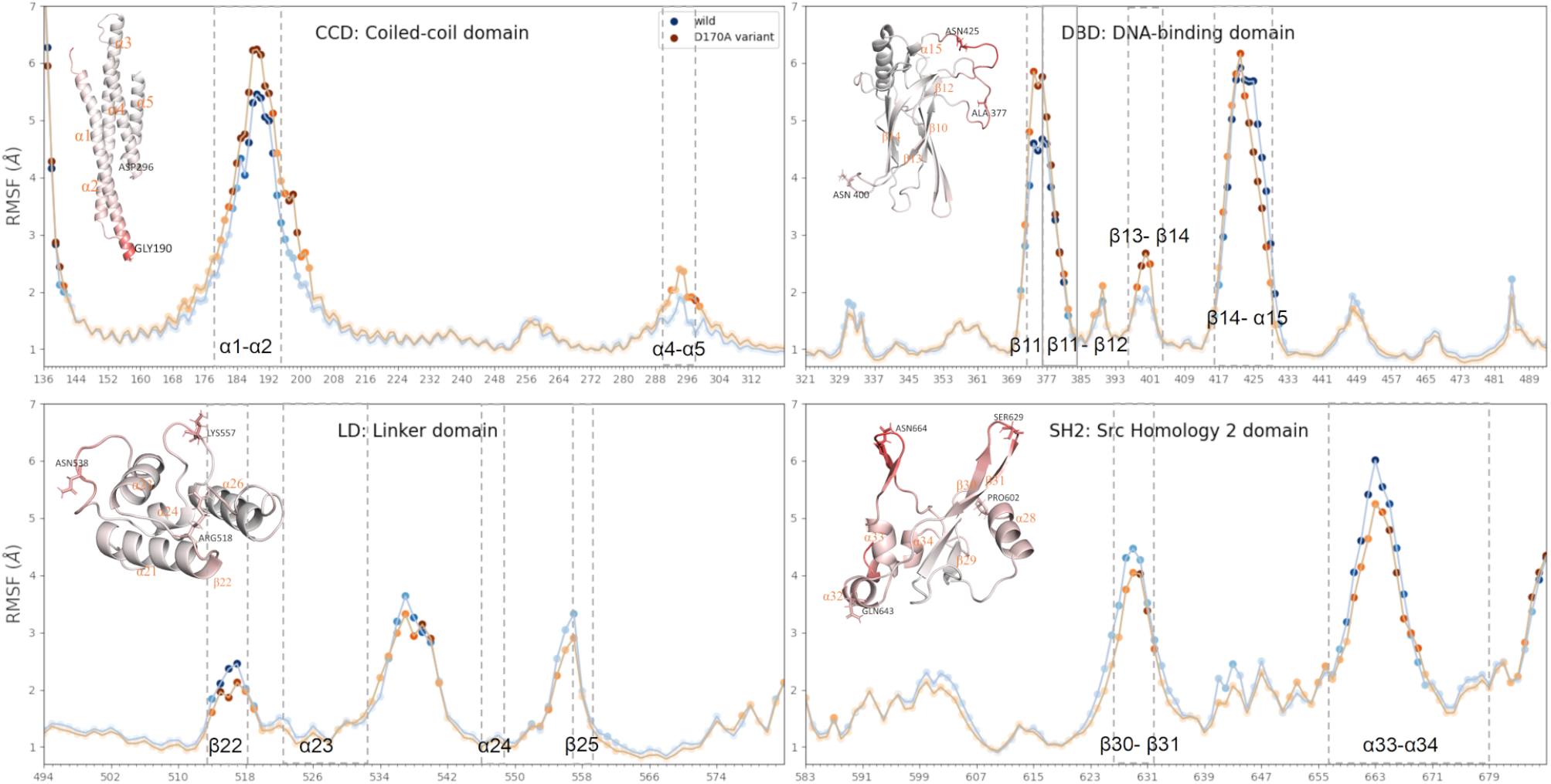
The mean RMSF of six replicates was plotted, with the RMSF values for each residue separated by domain. The wild type is plotted in blue and the D170A variant in orange, with the deepness of color indicating the stand deviation among the six replicates. Structures of each domain colored by wild type RMSF values are shown (low RMSF values in white to high RMSF values in red).

### Conformational differences in the pY+3 binding pocket correlate to the binding affinity decrease from wild type to D170A variant

The SH2 domain mediates binding of kinase-complexes to unphosphorylated STAT3, directing the phosphorylation of Y705 at TAD. Furthermore, the SH2 domain also provides the interface for dimerization of the phosphorylated TAD to form a functional pSTAT3 homodimer. Thus, any changes to the conformation of the specificity-determining region (pY+3 pocket) as well as the binding region of phosphorylated TAD (pY pocket) provides a key regulatory modification to STAT3 behavior. To describe the binding pocket conformations of SH2 domain, the pair center of mass (COM) distance matrix of key residues (Figure 4D) for both the pY and pY+3 pockets were calculated, separately. Principal component analysis (PCA) was employed to project the high dimensionality of pair COM distances into a 2D plane. The conformational space of both pockets for wild type and D170A variant is shown in Figure 4A (note that the principal components are determined from the combined wild type and mutant trajectories). The first two principal components contribute 62% of the total variation (Figure S2A), encapsulating the majority of the conformational space in just two dimensions.

**Figure 4:**
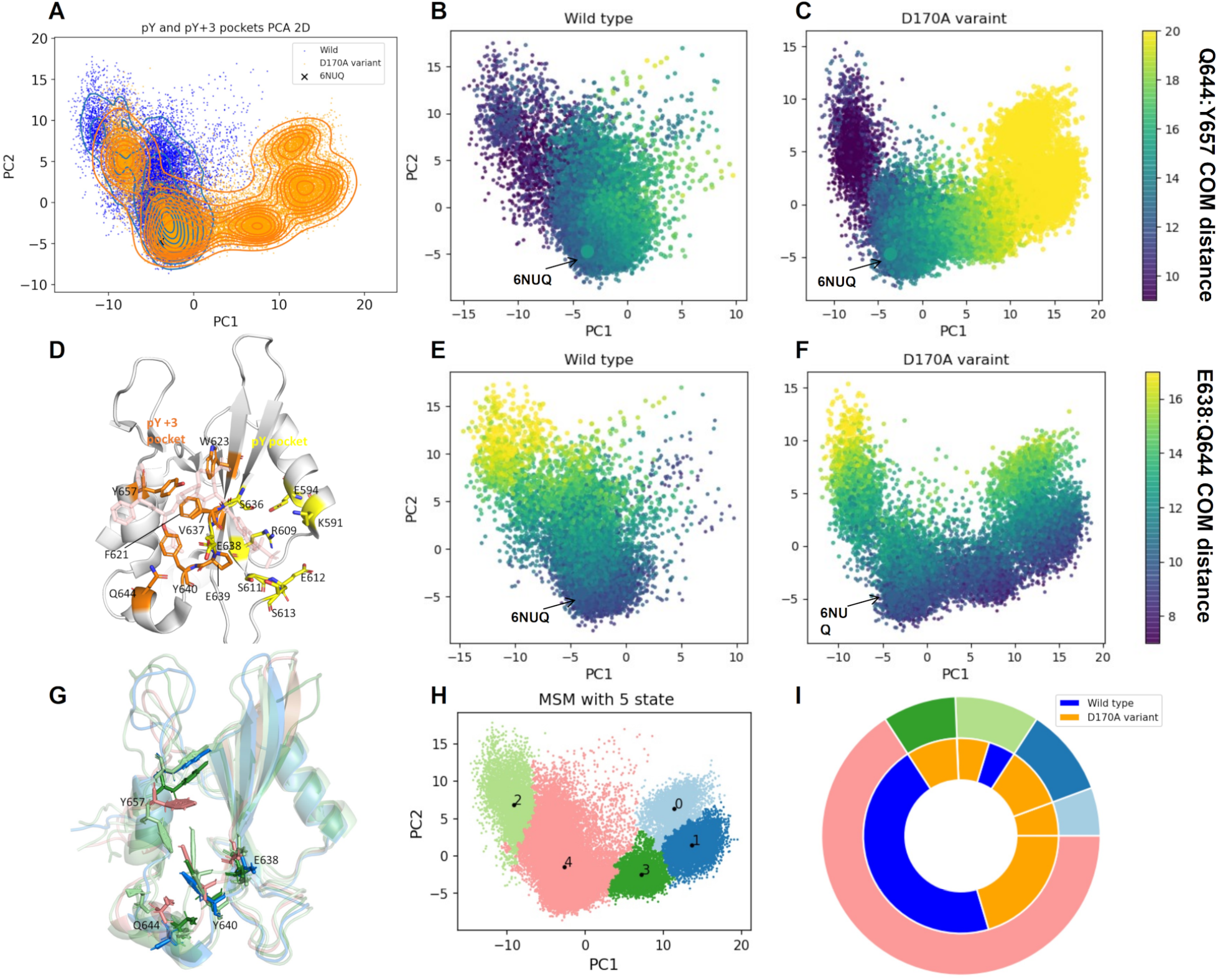
Conformational analysis of SH2 domain binding pockets. **(A)** PCA of both the pY and pY+3 pockets in the wild type (blue) and D170A mutant (orange). The contour lines show the density of recorded frames in each region, and the crystal structure (PDB ID: 6NUQ) is marked for reference. **(D)** SH2 domain bound to SI109 (light pink) to demonstrate binding mode of ligand to the pY and pY+3 pocket (6NUQ, Bai et al. ^11^). The pY pocket (yellow) and pY+3 pocket (orange) are shown with key residues included in the COM pair distances are shown as sticks. **(B**,**C)** PCAs for wild type and D170A variant, respectively, colored by the COM distance from Q644 to Y657. **(E**,**F)** PCAs for wild type and D170A variant, respectively, colored by the COM distance from Q644 to E638. **(G)** The averaged structures for each macro-state, residues were shown as stick. **(H)** pY and pY+3 pockets PCA 2D plane colored by different macro-states. **(I)** Nested pie charts for different macrostate by different system, outer circle was colored by different macro-state, inner circle was colored by different system

In the combined PCA plot, the conformational space of the D170A variant partially overlaps with the wild type, while also exploring distinct novel conformations (Figure 4A). The additional conformational space explored by the D170A variant occurs predominately along the PC1 axis. The ten largest coefficients of the first two PCs highlight the pair of residues with largest relative motions across all the trajectories (Figure S2C). The motion of Y657 and Q644 yields the largest coefficients across both PC1 and PC2, underlining their conformational importance. The Q644–Y657 pair has the largest variance along PC1(Figure 4B and C) while the E638–Q644 pair has the largest variance along PC2 (Figure 4E and F).

Comparing Figures 4 B, C, E and F with Figure 4A, we see that the E638–Q644 pair has a slightly higher variance in the wild type whereas the Q644–Y657 motion, particularly above ∼ 15, is predominately a feature of the D170A variant. Additionally, the D170A variant explores somewhat shorter Q644–Y657 distances than in the wild type. Most notably, all of the residues with highest variance occur in pY+3 pocket rather than the pY pocket. This result is in contrast with observation by Zhang et. al., where the authors show that D170A reduces binding affinity of ligands targeting the pY pocket. On the other hand, the binding mode analysis performed by Dhanik et. al. shows that the binding affinity of a ligand is directly correlated to additional interactions in the pY+3 pocket. High variance in the pY+3 pocket could potentially explain the reduction of binding affinity observed by Zhang et. al. without completely disrupting the pY pocket. Besides, Zhang et. al. observed that the truncation of STAT3 to exclude α1 helix is fully capable of DNA binding upon tyrosine phosphorylation by Src kinase in vitro, suggesting the preservation of a functional conformation of the pY pocket. This agrees with our observation of a stable pY pocket in the D170A variant.

To further investigate key differences in the overall structure, a combination of Markov State Modeling and Perron Cluster Cluster Analysis was applied to cluster transient conformations into kinetically meta-stable macro-states. Clustering into five macro-states (0–4) was applied to the combined wild type and mutant conformations (Figure 4H).

Both the D170A variant and wild type are well-represented within macro-states 2 and 4, while macro-states 0, 1 and 3 were uniquely explored by the D170A variant (Figure 4I). The average structures of each macro-state were calculated and are shown in Figure 4G. Distinct conformations of Q644 and Y657 are observed for each of the macro-states, in agreement with the PCA data. The pY+3 pocket is blocked in macro-state 2 by Y657 and Y640, both of them pointing towards the pocket. Conversely, in macro-state 0, 1, and 3, Y657 points away from the pY+3 pocket leading to an open conformation. The shared conformational states between wild type and D170A variant consists of a viable functional state of STAT3, however, D170A variant has reduced occupancy at those conformational states (Figure 4I), thus leading to a differentiated function relative to the wild type.

### Allosteric inactivation of SH2 pY+3 occurs via translation of motion through a rigid core

To explore the correlation between SH2 domain conformational changes and CCD conformations, CCD was first characterized via PCA analysis of all pair Cα distances. However, as the CCD domain consists of rigid helices, pair Cα distances were not able to characterize the differences between the determined macro-states, as indicated by the low variance contribution of each pair Cα distance (Figure S3A and C). Only macro-state 3 showed significant differences, which originate from kinked conformations of α1 (Figure S3B).

While the rigidity of the α helices diminished the utility of PCA analysis, it also demonstrates that the dynamics of the helices are highly correlated (Figure 3). This result led us to the hypothesis that the long helical arms of CCD may act as rigid levers, where a slight helical tilt causes significant changes in inter-domain interaction sites. The hypothesis was tested by comparing corresponding helical tilt at the CCD of different macro-states identified in Conformational differences in the pY+3 binding pocket correlate to the binding affinity decrease from wild type to D170A variant for the pY and pY+3 pockets. Significant differences in global helical tilt for each of the macro-states of the SH2 domain is observed in α3 showing functional correlative differences among macro-states (Figure 5H). The α3 helix also interfaces with most of the other domains through its C-terminal helical turn. Residues at this interface could transmit motion through interacting residues from other domains.

**Figure 5:**
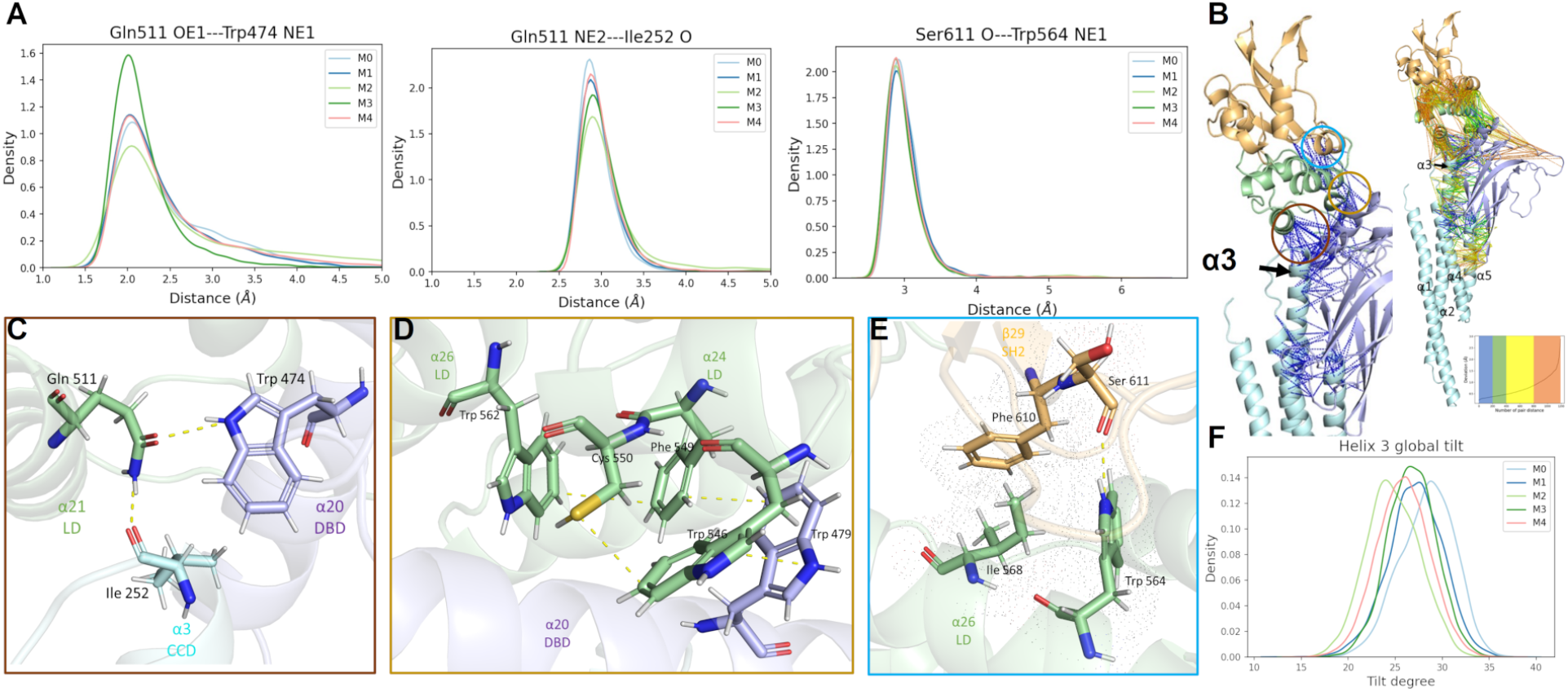
Rigid core analysis. **(A)** Key hydrogen bonds distance distributions. **(B)** Interdomain pair Cα distances shown as dashed lines and colored by normalized standard deviation across all trajectories. A close up of regions in colored circles are shown in C-E with respective colored borders. **(C)** Conserved hydrogen bond network between CCD, LD, and DBD. **(D)** Conserved π–π interaction network between LD and DBD. **(E)** Conserved hydrogen bond network as well as hydrophobic interactions between LD and SH2. **(F)** Distribution of global helical tilt of α3 in CCD for each macro-state.

### Conserved C*α* pair distances

The rigidity transmission was oberserved in other proteins (Sljoka ^23^, Ye et al. ^24^). Thus we further hypothesized that this motion is transmitted allosterically to SH2 via a “rigid core”, that is, an interlocking sequence of conserved interactions which function as a sort of molecular machine. The existence of such an interaction network is demonstrated in Figure 5B, where the inter-domain pair Cα distances are plotted and colored according to standard deviation values computed across all trajectories. There is a rigid backbone through the protein from CCD, LD, DBD, and finally to SH2 (Figure 5B) which is highly conserved during dynamical motion of the protein before and after mutation at D170. α3, α20, α21 compose the first section of the rigid core between CCD, DBD, and LD (Figure 5C), which could convey the dynamics of CCD into this highly rigid region. Upon close inspection, the three helices α3, α20, and α21 are locked via hydrogen bond network between I252, Q511 and W474 (Figure 5A,C), such that any rotation of these residues results in a corresponding reorientation of the helices to preserve the hydrogen bond network. We additionally find strongly conserved inter-domain interactions between the DBD and LD (Figure 5D), as well as between the LD and SH2 (Figure 5E), which complete the rigid core. Subtle changes in the CCD configurations can thus convey movement from α3 through α21 (in LD) and α20 (in DBD) to α26 and α24 (in LD), and finally leading to allosteric modification of SH2 via β29.

The conformation analysis presented above demonstrates that D170A mutation leads to changes in the orientation of the CCD α helices, which could then lead to allosteric regulation of SH2 domain conformations though a network of conserved interactions. Rigid body analysis shows that these interactions consist of hydrogen bond, π–π, and hydrophobic networks that strongly correlate motions of different domains. However, this analysis does not highlight the allosteric path that differs between the wild type and D170A variant nor provide evidence of a dynamical correlation between CCD and SH2 conformation through this pathway. To further elucidate the allostery pathway and show dynamical correlation, we employ both a REDAN analysis and and analysis of differences in the global hydrogen bond network between kinetic macro-states.

### REDAN

While a rigid backbone provides a potential pathway for signal propagation, the cumulative long-range allosteric effect is realized through short range interactions and subtle allosteric changes which must occur in concert. To identify a detailed sequence of short range interactions, REDAN analysis was employed as a means to identify residue pairs that are responsive to allosteric perturbation, followed by shortest path analysis using Dijkstra’s algorithm. Using REDAN, subtle yet highly correlated differences in the allosteric network between D170A and the wild type can be resolved allowing us to propose a concrete signal transduction network from CCD to SH2.

From the conformational analysis, Y657 was identified as the residue with highest average difference between different macro-states explored by the SH2 domain. Thus, D/A170 was selected as the starting point and Y657 was selected as the end point for the analysis. The most structurally-relevant pathway from effector residue (D/A170) to regulatory site (Y657) was identified by REDAN and is shown in Figure 6A,B. The pathway originates from the CCD, through the LD and to the SH2 domain, bypassing DBD (although it passes nearby α20 which was identified as a component of the rigid core). The residue pairs that connect domains are of most interest, and their distance distributions are shown in Figure S4 C, D, and E. The average structures of each macro-state were calculated to structurally verify correlated motions in secondary structure identified by the REDAN (Figure 6C).

**Figure 6:**
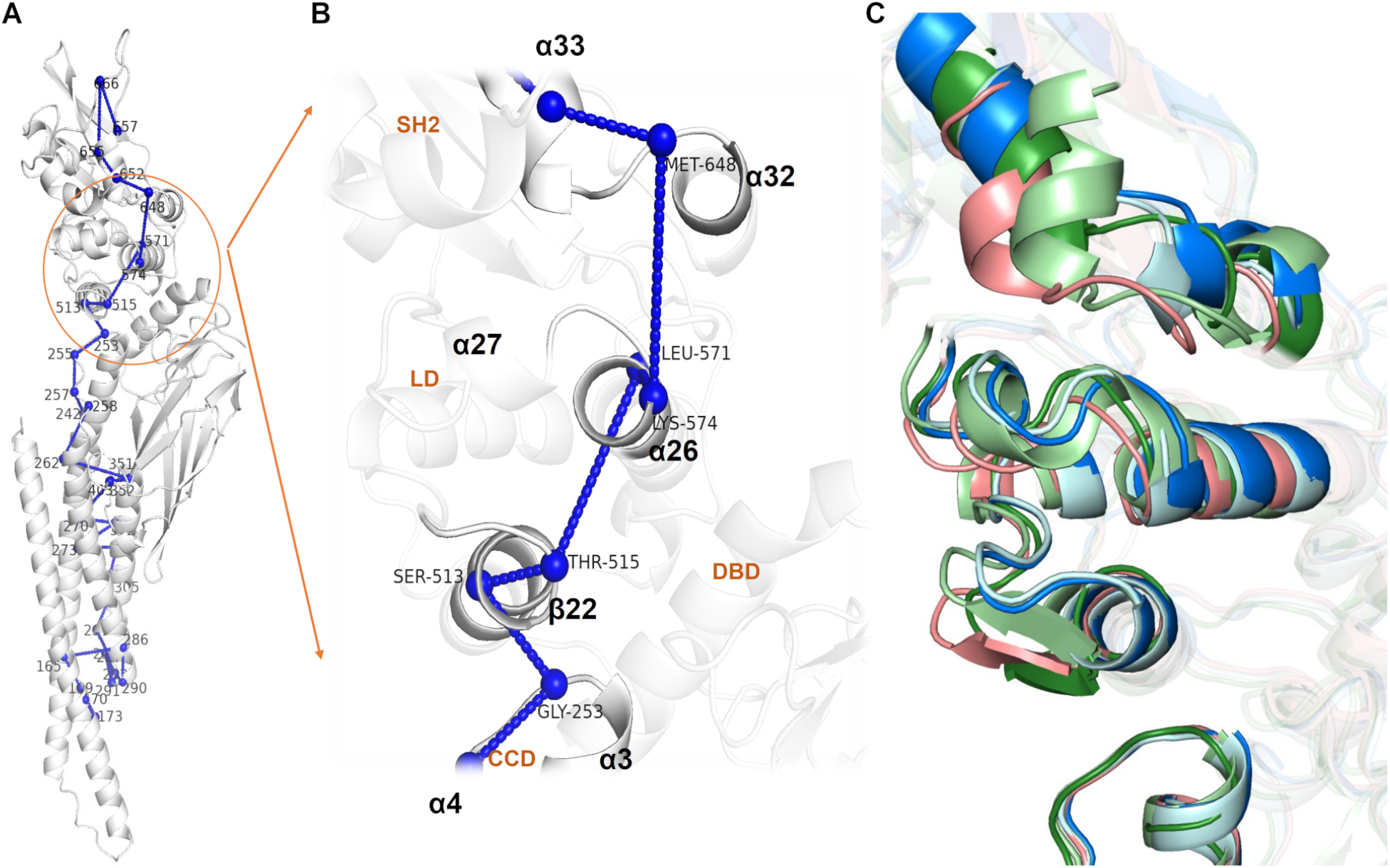
Proposed allosteric path from D/A170 to Y657 obtained from REDAN analysis and its structural details. **(A)** Residues involved in the allosteric path from CCD to LD and to SH2 domain. The raw path can be found in (Figure S4). **(B)** A close up view of signal transduction from CCD domain to SH2 domain. The residues identified by REDAN show a path through α3 in CCD to α21 and β22 in the LD domain and finally to α32 and α33 in the SH2 domain, bypassing DBD. **(C)** Average structures from all the macro-states show significant reorganization in this interface (colors as in 4 H). β sheet configuration (adjusted in PyMol) of macro-states 2, 3, and 4 are shown to highlight rearrangement of β22 based on Ramachandran Dihedral for β-sheets of residues 514 to 517.

The secondary structure designated β22 (Figure 1B) can be observed to dynamically shift between a β sheet and α helix in macro-states 2, 3, and 4, while in macro-state 0 and 1 the α helix is stabilized (Figure S4F). β22 extends the α21 helix and reorganizes the loop between β22-α23. It is worth noting that residues 514 to 517 are annotated as a β sheet in the UniProt database (Table S1), while these residues form an α helix in the reference structure used in this study (PDB ID: 6TLC). This result is not necessarily incongruous with the database designation as multiple configurations of the secondary structure are observed experimentally. Stabilization of β22 as an extension of the α21 helix in macro-states 0 and 1 stabilizes extension and a shift of the α26 helix. The REDAN analysis suggests that further interaction between α26 and α32 then positions the α32-α33 loop such that α33 adopts an extended conformation without close contact to the α32-α33 loop. On the other hand, the structural conformation analysis above suggests that α26 may more indirectly affect α32 via hydrogen bonding and hydrophobic interaction with β29.

### Hydrogen bond network

The formation and breaking of hydrogen bonds plays an important role in stability of secondary structures and conformational variability of tertiary structures of a protein. To complement structural insights and REDAN analysis, the differential rate (preponderance) of hydrogen bond occurrence between different macro-states was compared. Hydrogen bonds were identified using Baker-Hubbard hydrogen bonding analysis, and then the differential hydrogen bond rate was calculated between macro-states 0, 1, 2, and 3, and the wild type-dominant macro-state 4. Macro-state 4 was considered as the primary active state as the wild type and D170A share high occupancy rates at this state. Note that Baker-Hubbard hydrogen bonding analysis algorithm classifies salt-bridge formation between amine and carboxylic acid as a hydrogen bond as well, thus the formation and breaking of salt bridges were also considered in the analysis.

Consistent with the observation from REDAN analysis, β22 has large changes in the hydrogen bond between the native (macro-state 4) and allosteric (macro-states 0 and 1) configurations. In macro-states 0 and 1, hydrogen bonds formation in β22 stabilizes the alpha helix, while the other macro-states alternate between an α helix and a β sheet. Moreover, in macro-state 3, rearrangement causes new a hydrogen bond to form between β22 and α3. These changes highlight key differences between the wild type and D170A variant as only the D170A variant occupies macro-states 0, 1 and 3. These macro-states also show decoupling of LD to DBD along with alteration of interactions between LD and SH2 (Figure 8). First, a salt bridge between D566 (LD) and R335 (DBD) is lost in macro-states 0 and 1 (Figure 8B,C). The loss of this interaction allows for the shift of α26 observed in the macro-states (Figure 6C). Second, a new hydrogen bond is formed between I576 (LD, α26–α27 loop) and N646 (SH2, α32) in macro-state 0 (Figure 8B) and a salt bridge is formed between D570 (LD, α26) and K642 (SH2, α32) in macro-state 3 (Figure 8C). Third, loss of hydrogen bonds of E652 and/or I653 (LD, α33) with S649 (LD, α32–α33 loop) in macro-states 0, 1, and 3 (8A–C) contributes to increased flexibility of the α33 helix, also as observed in Figure 6C.

**Figure 7:**
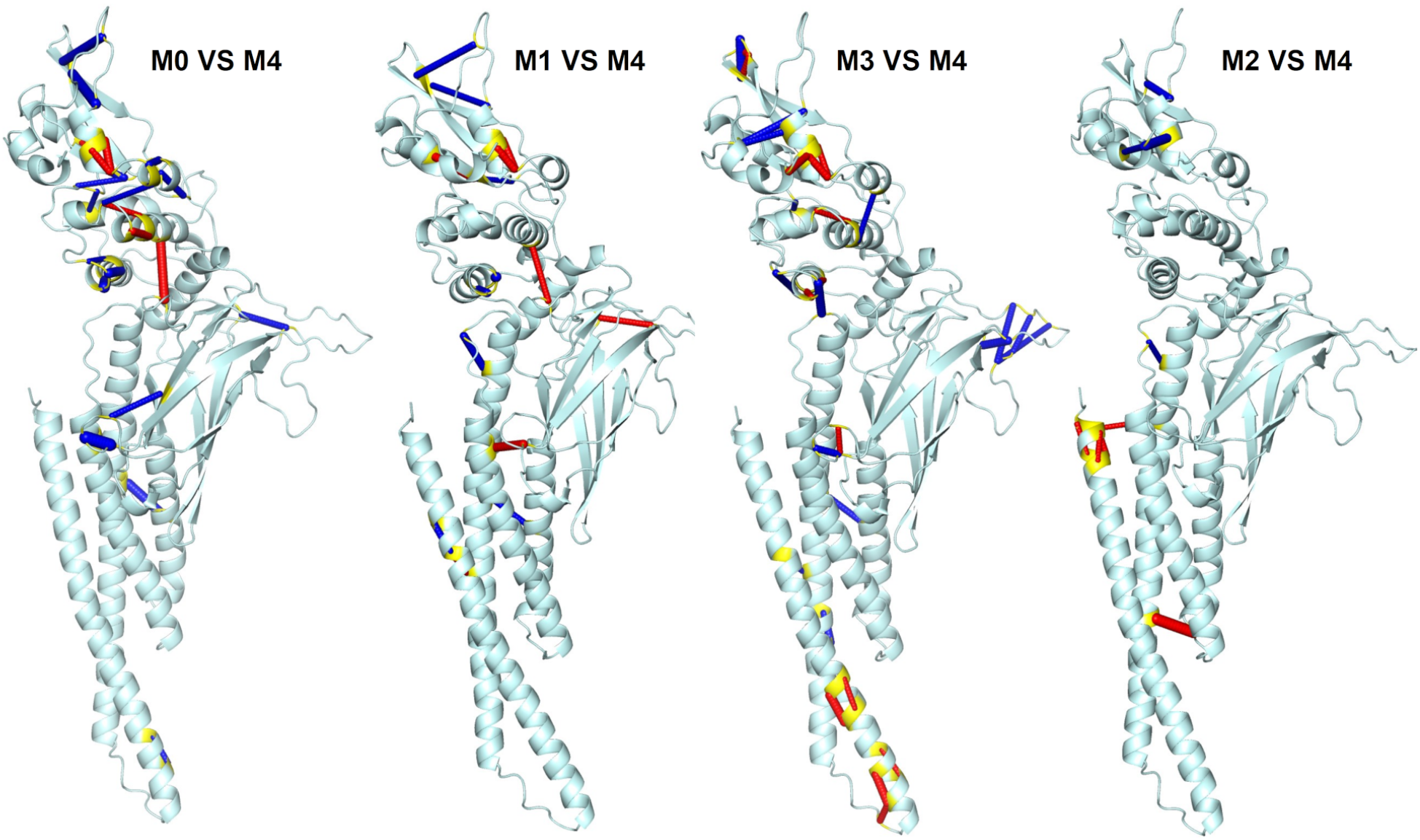
The differential rate of pair residue hydrogen bonds for macro-states 0–3 compared to macro-state 4. Using the left-most structure as an example, blue links indicate that a hydrogen bond is present in at least 50% more frames in macro-state 0 compared to macro-state 4, while red links indicate a rate at least 50% lower. Terminal residues are colored in yellow. While significant changes in hydrogen bonding are evident across the protein, macro-states 0, 1, and 3 clearly indicate large changes in the pY+3 pocket, as well as consistent hydrogen-bonding changes linking SH2 and LD domains.

**Figure 8:**
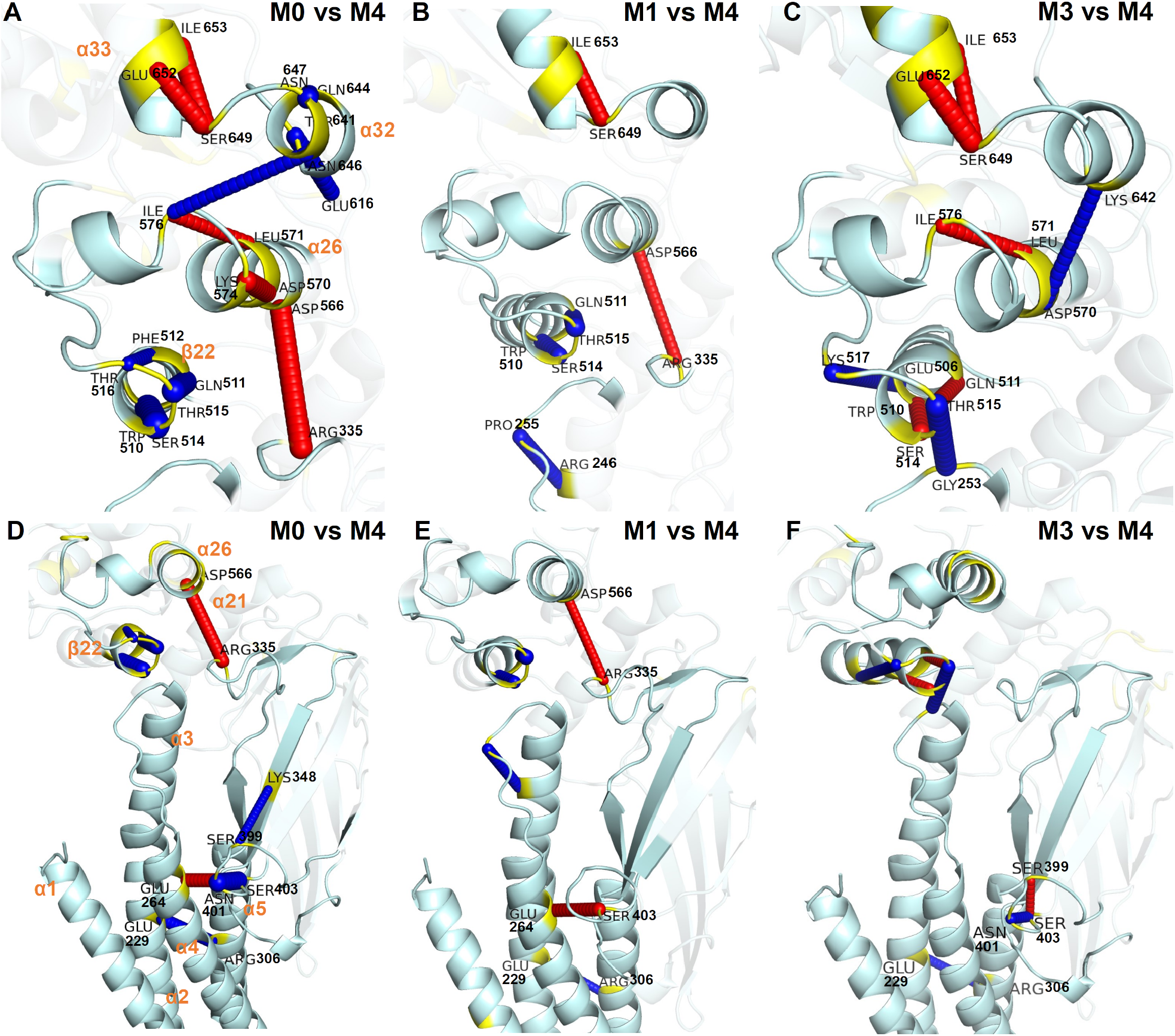
A close up view of the hydrogen bond network changes between macro-states 0–3 and macro-state 4. **(A–C)** Differential rate of pair residue hydrogen bonds in the LD and SH2 domains. **(D–F)** Differential rate of pair residue hydrogen bonds in the CCD and LD domains.

Focusing on the upper part of CCD (Figure 8D–F), it is clear that α1 interacts directly with α2 and α4 through a large interaction surface. α2 and α3 form a contiguous helix interrupted by a kink in the helix at residue 278, thus perturbation of α2 is structurally coupled with α3. A salt bridge between E229 (α2) and R306 (α5) has a high appearance rate in macro-states 0, 1, and 3 compared to macro-state 4. This salt bridge provides a means to strongly correlate the motion of α2 to α5 and mitigate perturbations by α1. These interaction alter the overall helical tilt of α3 which is translated to the rigid core interface. Moreover, I252 resides at the terminal end of α3 which forms strong hydrogen bond with Q511 (LD) identified in the rigid core analysis (Figure 5C). Q511 also forms a strong hydrogen bond with W474 (DBD). This network of hydrogen bonds allows a strong correlated motion between α3 in CCD and α20/α21 in the LD domain. I252 is also adjacent to E253 which was identified by REDAN analysis as part of the allosteric pathway thus providing a chemical basis for signal transduction from CCD to SH2 via the LD domain.

## Discussion

Significant differences in conformational space of the SH2 domain binding pocket (pY+3) between the wild type and D170A mutant were observed. The D170A variant explores and extends conformational space of SH2 domain, specifically with significant changes in opening of the pY+3 pocket. The hydrophobic environment formed by I659, W623 and F621 in pY+3 were previously shown to assist in binding of target peptide. Changes in the hydrophobic environment by increasing either hydrophobicity or aromaticity leads to hyperactivation, while introduction of polarity and reduction of hydrophobicity and aromaticity diminishes STAT3 function. (de Araujo et al. ^25^). Furthermore, studies suggest that the side-chain Y657 interaction is important for stabilizing the ligand-protein complex (Dhanik et al. ^15^, Mandal et al. ^16^, McMurray ^17^). Thus, the diminished binding affinity can be attributed to increased motions in the structures surrounding the pY+3 pocket in D170A. While in principle, translation of motion through the rigid core could affect the conformation of the primary pY binding pocket, only the pY+3 pocket shows differential motions in D170A compared to the wild type. Thus, our simulations support the conclusion that mutation of the D170 residue affects an inactivation of the protein via an allosteric mechanism resulting in conformational changes primarily in the specificity determining pY+3 pocket.

The observed differences in the SH2 conformational space allow us to further characterize the mechanism of this allosteric effect. The α3 helix of the CCD domain correlates functionally with SH2 conformations, as evidenced by the differing global tilt angles observed for different macro-state clusters. Given its positioning, it likely communicates changes in the CCD to the LD. Using rigid body analysis, a potential pathway from CCD through LD, and finally to SH2 was identified, wherein conserved hydrogen bond networks and other strong interactions firmly link a series of secondary structures (primarily α3, α20, α21, and α24). While further analysis supports this proposed mechanism (vide infra), controlled mutagenesis of these key residues could also provide experimental evidence for the importance of these interactions to the D170A allosteric pathway. The lack of a significant increase (or decrease) in dynamic motion and overall flexibility in D170A compared to the wild type (as seen in the RMSF analysis) also supports a sequence of interactions between rigid bodies as the main allosteric mechanism.

The specific allosteric pathway was further elucidated via REDAN and differential hydrogen bonding analysis. These analyses both point to a very specific mechanism (structural details shown in Figure S5): 1) stronger interaction between α5 and α2 causes a tilt in the α2/α3 helix, 2) α3 tilt interfere the interaction between β22/α23 and α26/α27. In macrostate 0 and 1, the interference breaks the salt bridge between D566 and R335 and caused α26 shift away β22. While in macrostate 3, α26 shift toward β22, which cause a steric clash between Lys 573 and β22. 3) in turn, the movement of α26 and α26/α27 causes breakage of the hydrogen bonds between α33 and the α32-α33 loop, 4) α33 extends significantly and alters the conformation of the pY+3 pocket. We also identified conserved interactions between α26 and β29 in SH2 which may provide further coupling.

Previously, the role of α26 in allosteric communication has been identified: nuclear magnetic resonance (NMR) studies showed that mutation of I568F was able to induce a chemical shift perturbation in SH2, DBD, and CCD (Namanja et al. ^26^). Besides, D566A, D570A and D570K mutants showed profound negative effects on transcription, and also unexpectedly tyrosine phosphorylation even before IL6 induction. (Mertens et al. ^27^, Yang et al. ^28^). Additionally, previous study shows similar allostery pathways in STAT5, however rigid core was not explored (Langenfeld et al. ^29^). STAT family of proteins are highly similar in primary, secondary and tertiary structures and the similarity in allosteric pathways of STAT3 and STAT5 can be used to posit that STAT family of protein have a rigid core that couples allostery between their domains.

These results present us with novel ways of regulating the CCD domain whereby ligand or peptide interaction with CCD can significantly alter the helical tilt of α3 which is transmitted to the SH2 domain via the identified allosteric pathway. It is unlikely that different effectors will have identical mechanisms within CCD, as the local perturbation caused by point mutation, small molecule binding, peptide binding, etc. is radically different. However, our analysis supports the conclusion that any perturbation resulting in a change of α3 tilt should result in a similar outcome due to the strong and highly concerted motions along the proposed allosteric pathway. Potentially, alteration of helical tilt could also result in tighter binding of target peptides to the SH2 domain while, as observed here, helix-helix interactions in CCD can also promote structural changes in SH2 domain leading to reduced affinity of SH2 binding. Drugs designed to specifically alter the motions of the CCD helices would yield valuable insight towards validation of proposed mechanism. Since the pY+3 pocket regulates the affinity of peptide binding, rather than being the catalytic active site, assays developed to probe allosteric regulation of SH2 via CCD should consider the identity of peptides used to target the SH2 domain in addition to overall activity.

Finally, the methods used here to identify the rigid core mechanism, specifically a combination of structural and conformation analysis (dimensionality reduction, functional clustering, and conserved Cα pair distances) with dynamical and correlative analyses (REDAN and differential hydrogen bond analysis) should allow for identification of other potential effectors targeting CCD, and more widely, in identifying similar allosteric pathways in a number of (semi-)rigid proteins.

## Methods

### Initial structure

The monomer STAT3 with peptide MS3-6 complex (PDB ID: 6TLE) structure was used as the template type for this study. The missing residues were modeled using Chimera (Pettersen et al. ^30^). The apo structure was created by directly deleting the MS3-6 peptide, and then the apo type was subjected to mutation using the PyMol Mutagenesis Wizard (Schrödinger, LLC ^31^) to generate the D170A variant.

### Molecular dynamics simulation

For each system, a rectangular periodic water box of TIP3P waters was used with a minimum distance of 10 between the box boundary and the protein to avoid image interactions. To balance charge and provide realistic salinity, 0.15M sodium and chloride ions were added. NAMD 2.13 (Phillips et al. ^32^) with the CHARMM 36 force field (Best et al. ^33^) was used for energy minimization and molecular dynamics (MD) simulations. Initially, the simulation systems were subjected to 5000 steps of energy minimization to remove bad contacts and clashes. Then, systems were heated from 0 K to 300 K, heating 50 K every 200 ps, and then from 300 K to 310 K in 200 ps, with 10 ns isothermal-isobaric ensemble (NPT) short equilibration. Subsequently, six replicas of 600 ns canonical ensemble (NVT) MD simulations at 310 K were conducted. The first 100 ns simulations were discarded as equilibration and the following 500 ns for each replica, 3ms in total, was used for further analysis. The SHAKE algorithm was applied to all bonds containing hydrogen atoms. The electrostatic interaction was evaluated by the particle-mesh Ewald method, and Lennard-Jones interactions were evaluated using 10 as a cutoff.

### RMSD analysis

Root-mean-square deviation (RMSD) analysis shows the conformational dynamics over the trajectory and provides an insight into the flexibility of the system, and to identify variation within the conformational space of a reference structure. After optimal rigid-body superposition of the simulation frame to the reference structure, the RMSD is calculated is calculated as,

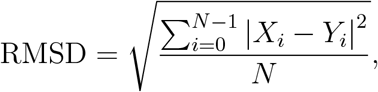

where *N* is the number of atoms, *X*_*i*_ is the coordinate vector for target atom *i*, and *Y*_*i*_ is the coordinate vector for reference atom *i*. The RMSD, RMSF, pair Cα distance, and hydrogen bond analysis were performed using MDTraj 1.9.3.40 (McGibbon et al. ^34^).

### RMSF analysis

Root-mean-square fluctuation (RMSF) allows us to probe average positional changes of each residue. RMSF measures the average deviation of a particle (and individual residue) over time from a reference position (typically the time-averaged position of the particle). First, the RMSF of a each atom *i* is calculated as,

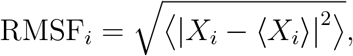

where *X*_*i*_ denotes the coordinate vector for atom *i*, and ⟨·⟩ is an average over all input frames within a replica. Within each residue *I*, the mass-weighted average of atomic fluctuations is calculated as,

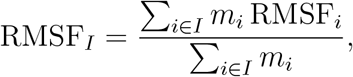

where *m*_*i*_ is the mass of atom *i*. Finally, the RMSF values of each residue are averaged over the six replicas.

### Principal components analysis

Linear Principal Components Analysis (PCA, Wold et al. ^35^) is used to transform high dimensional and often linearly-dependent data points into a low-dimensional space spanned by uncorrelated principle components. The first two principal components (PCs) were used in this work, yielding a two-dimensional reduction of the original data. Given high dimensional data represented in n (sample size) by m (variable size) matrix, the covariance of any two variables X and Y was calculated by,

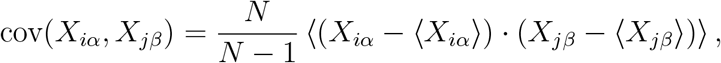

where *X*_*iα*_ is the *α* Cartesian component of the coordinate vector for atom *i*. The covariance matrix *C* is constructed as the pairwise covariance between all variables. The eigenvectors of *C* are the components of PCA, while the eigenvalues measure the contribution of each PC in the dataset. The eigenvectors also provide a mapping from the high-dimensional dataset to the low-dimensional PC space: the (PC1,PC2) coordinates for each input frame are given by multiplication of the original data set by the first two important eigenvectors. For a given PC, the importance of a feature (variable) is reflected by the absolute magnitude of the corresponding entry in the eigenvector. The PCA analysis was performed by Scikit-learn (Pedregosa et al. ^36^) implemented in Python.

### Markov state modeling and Perron cluster cluster analysis

Markov state models (MSMs, Husic and Pande ^37^, Bowman et al. ^38^) have shown great utility in modeling the transitions among functional states. It was used in this study to cluster conformations into kinetically meaningful macro-states. The conformational space was first discretized into *n* micro-states. Here, agglomerative clustering (Zepeda-Mendoza and Resendis-Antonio ^39^) was applied to divide the sampled conformations into 300 micro-states in the two-dimensional PCA coordinate system. Then, *C*_*ij*_(*τ*), the number of observed transitions from micro-state *i* to micro-state *j* at a lag time τ is calculated. Then the transition probability, *P*_*ij*_, from micro-state *i* to micro-state *j* can be estimated as *P*_*ij*_ ≈ (*C*_*ij*_ + *C*_*ji*_)/ Σ_*k*_ (*C*_*ik*_ + *C*_*ki*_). According to estimated relaxation timescale (Figure S2B), the count matrix and MSM transition probabilities converge beyond *τ* = 5 ns, which was chosen as the lag time.

Finally, Perron Cluster Cluster analysis (PCCA), implemented in the PyEmma package (Scherer et al. ^40^), was be used to coarse grain micro-states to “kinetically relevant” macrostates based on the well-sampled micro-state transition matrix. Structures that interconvert frequently were assumed to belong to the same functional metastable state (macro-state) (Pande et al. ^41^). Five macro-states were determined based on the band gap in the estimated relaxation timescale plot.

### Relative entropy-based dynamical allosteric network

The relative entropy-based dynamical allosteric network (REDAN) model (Zhou and Tao ^42^) was used to quantitatively characterize protein allosteric effects upon mutation. The difference between the distributions of the pair alpha carbon (Cα) distances of two residues upon perturbation is quantified by the perturbation relative entropy (PRE), which is the average relative entropy,

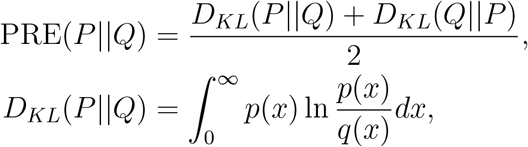

where *p*(*x*) is the distribution density for system *P* (before perturbation), and *q*(*x*) is the distribution density for system *Q* (after perturbation). High perturbation relative entropy values indicate that substantial allosteric effects are implied by the significantly different distance distribution of the residue pair.

Then, a weighted graph can be built based on the PRE matrix. Each node is represented by a Cα atom, and two nodes will be connected by an edge if the longest possible distance between them is less than 8. Each edge is weighted as 1/PRE. Since high PRE values indicates importance in the propagation of the structural changes in the protein, the pathway with the smallest overall weight implies the most structurally-relevant route and hence a possible allosteric pathway. Dijkstra’s algorithm was used to identify the shortest pathway.

### Alpha helix global tilt

The α helix global tilt angle was used to characterize the geometry of helices according to the procedure of Sugeta and Miyazawa (Sugeta and Miyazawa ^43^). After alignment to the crystal structure, the Cα positions of four “equivalent” residues, spaced equally apart, are taken as the reference positions. The differences of the vectors connecting subsequent Cα atoms, *C* and *C*^*′*^, lie perpendicular to the helix axis. Thus, the γ tilt angle is given as the angle between *C* × *C*^*′*^ and a reference axis chosen relative to the crystal structure frame. The MdAnalysis software was used to characterize the alpha helices (Bansal et al. ^44^).

## Acknowledgements

This work was supported in part by the National Science Foundation under Grant No. (1753167). TZ was supported by a fellowship from the SMU Department of Chemistry. All calculations were performed on the ManeFrame II supercomputing system at SMU.

