## Supplemental information for "Allostery in STAT3 Variant D170A is Mediated by a Rigid Core"

### **Supporting information**

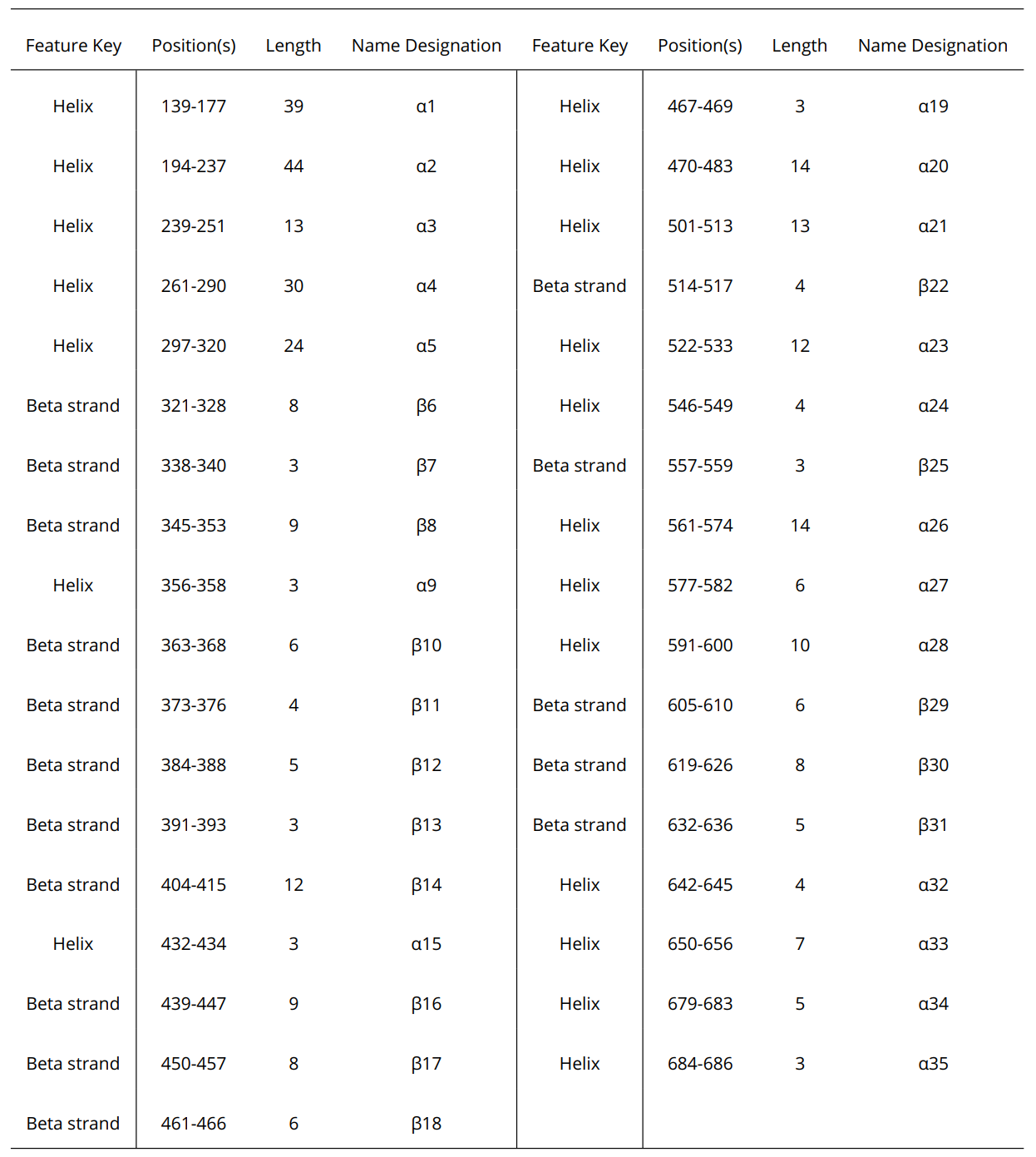

Table S1: Secondary Structure assigned by uniport database by compiling structure information from multiple x-ray crystal structures.

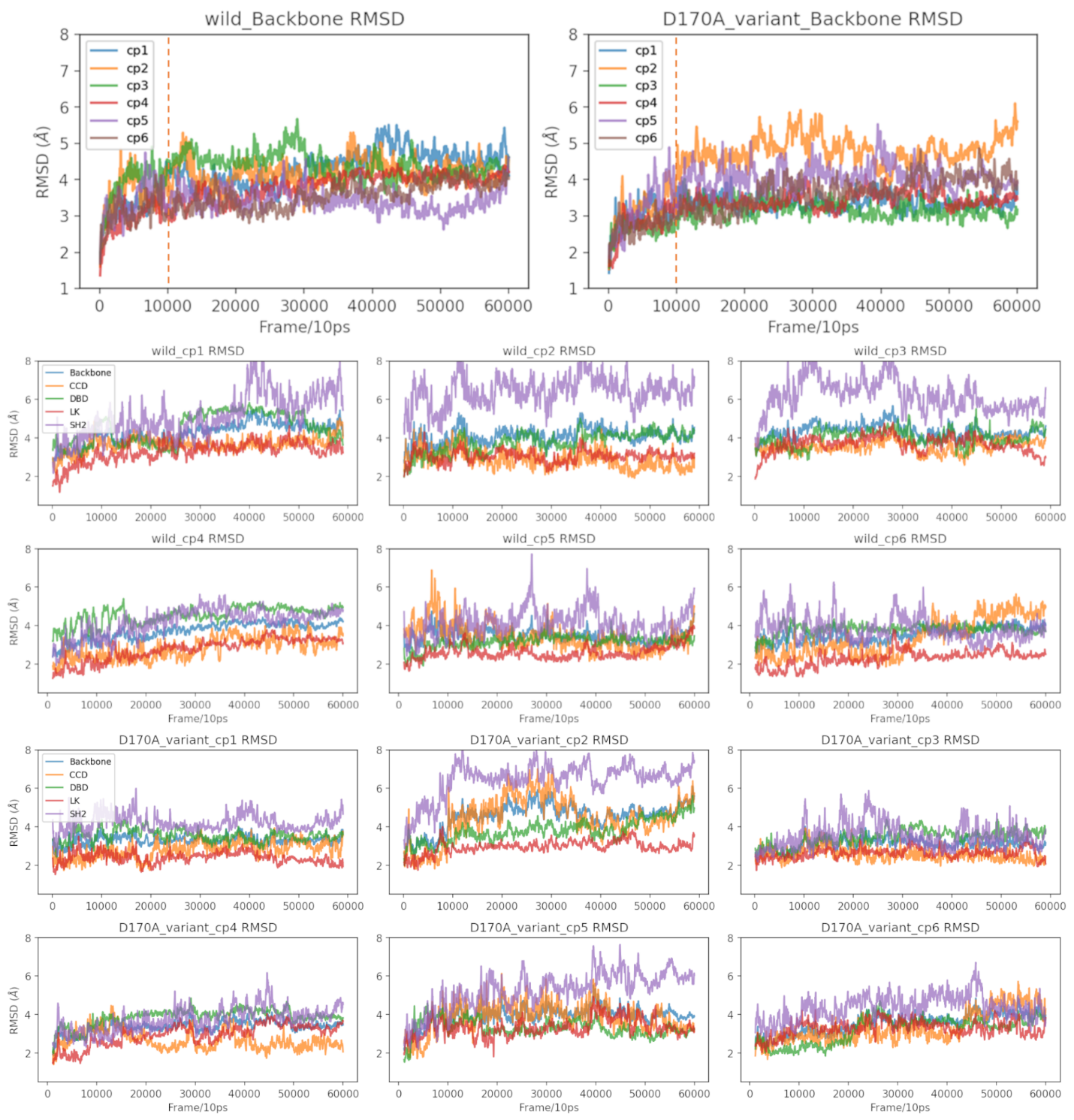

Figure S1: Root Mean Square Deviation (RMSD) of wild system and D170A variant system, each system has six replicas: cp1, cp2, cp3, cp4, cp5 and cp6. And the RMSD of each domain for each replica. Note: Rolling average of every 100 frames was plotted here for better visualization.

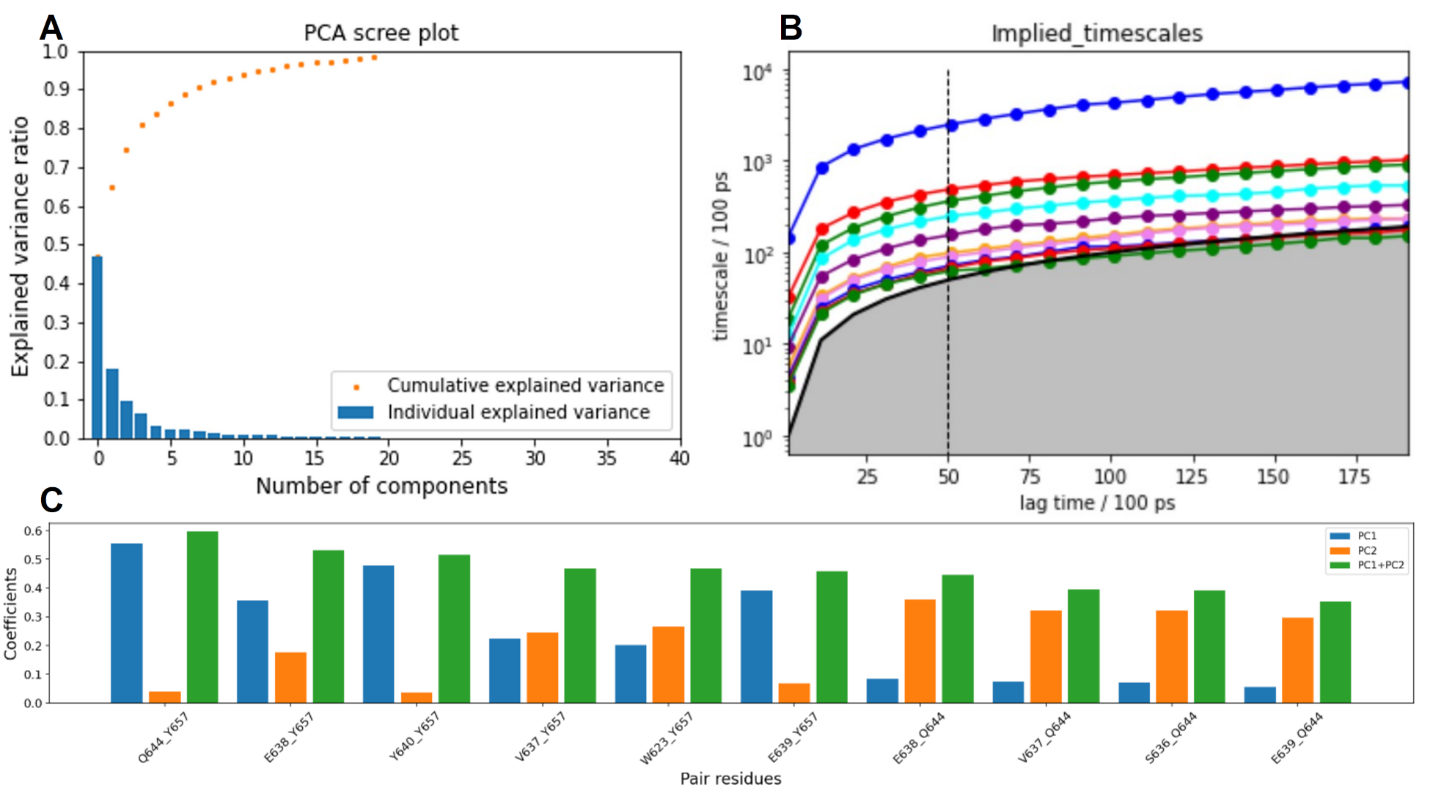

Figure S2: Additional information of PCA and MSM analysis. **(A)** PCA scree plot: dot shows the cumulative explained variance of the principal components; the bar chart represents the explained values per component. **(B)** Relaxation timescales of MSM for SH2 domain conformational space at different lag times. **(C)** The first ten features that contribute the first two PC the most.

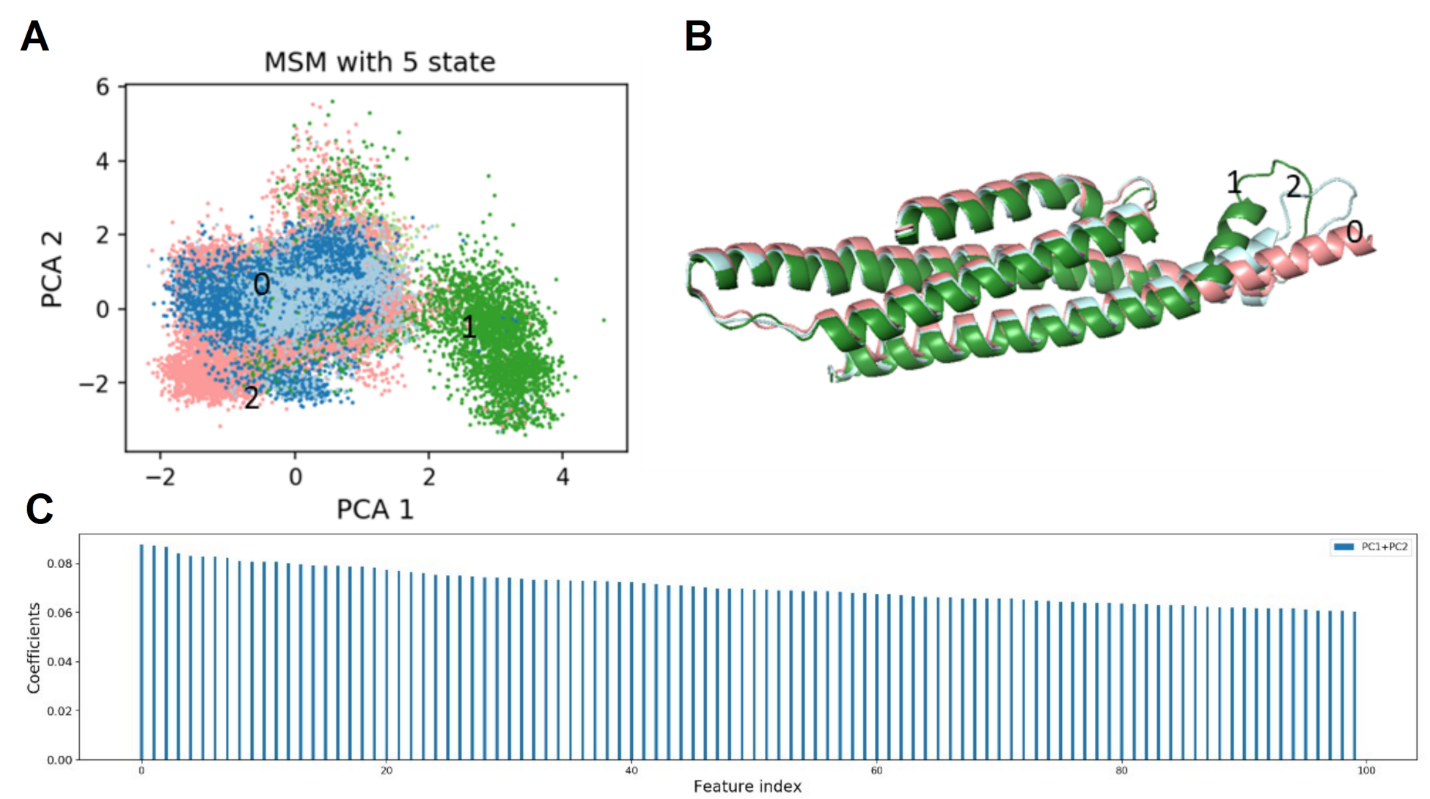

Figure S3: Characterization of CCD using Pair ca distances. **(A)** PCA 2D plane of CCD pair ca distances colored by Macrostate from SH2 domain results. **(B)** Represent structure of CCD corresponding to Figure A. **(C)** The coefficients of first 100 features that contribute the first two PC the most.

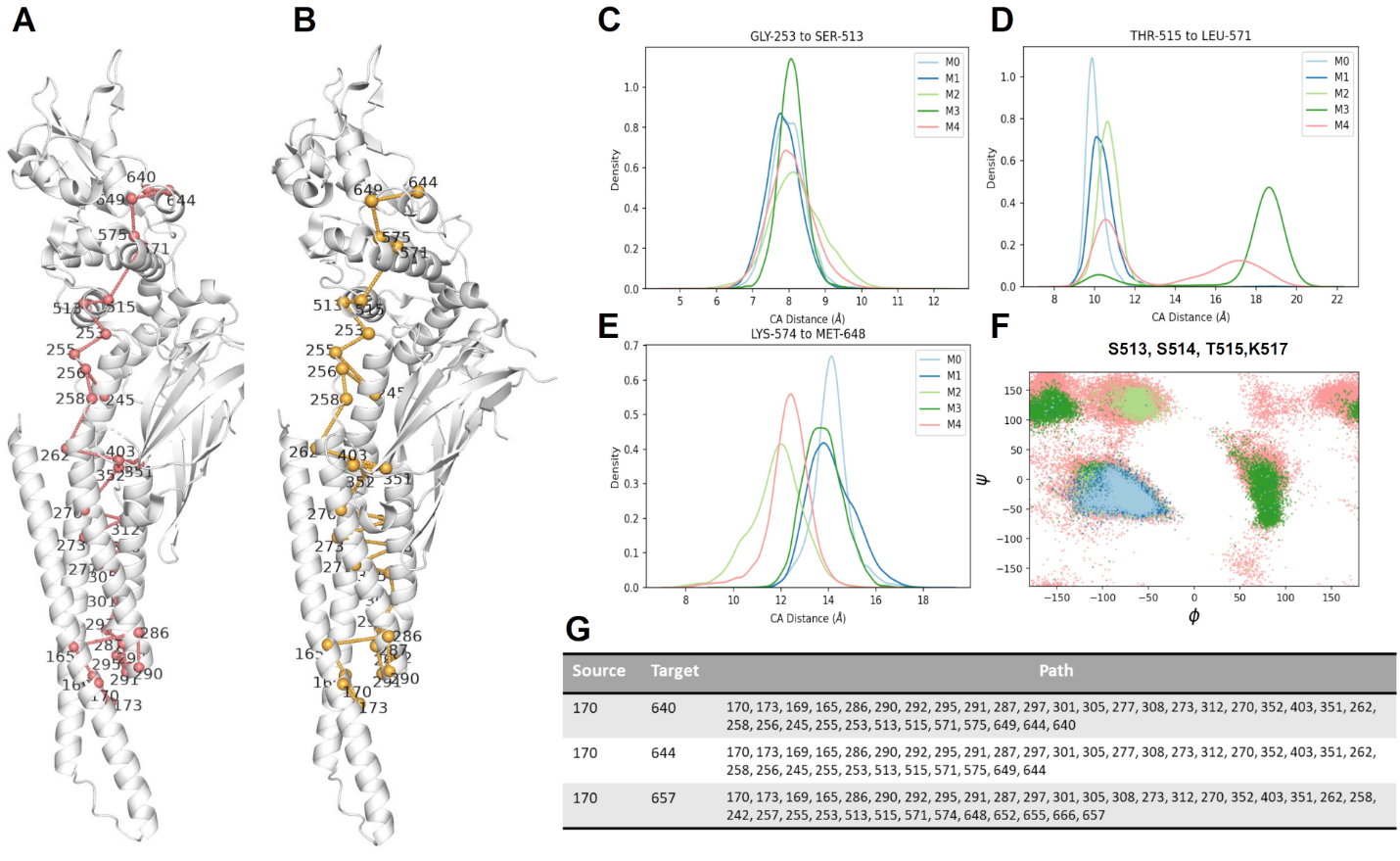

Figure S4: Additional REDAN analysis results **(A)** Proposed pathway from 170 to 640 shown in the protein structure; **(B)** Proposed pathway from 170 to 644; **(C,D,E)** Key pair residue distance ; **(F)** Ramachandran Dihedral for residue 513,514, 515 and 517; **(G)** Summary of proposed pathways from source residue 170 to target residue 640,644 and 657.

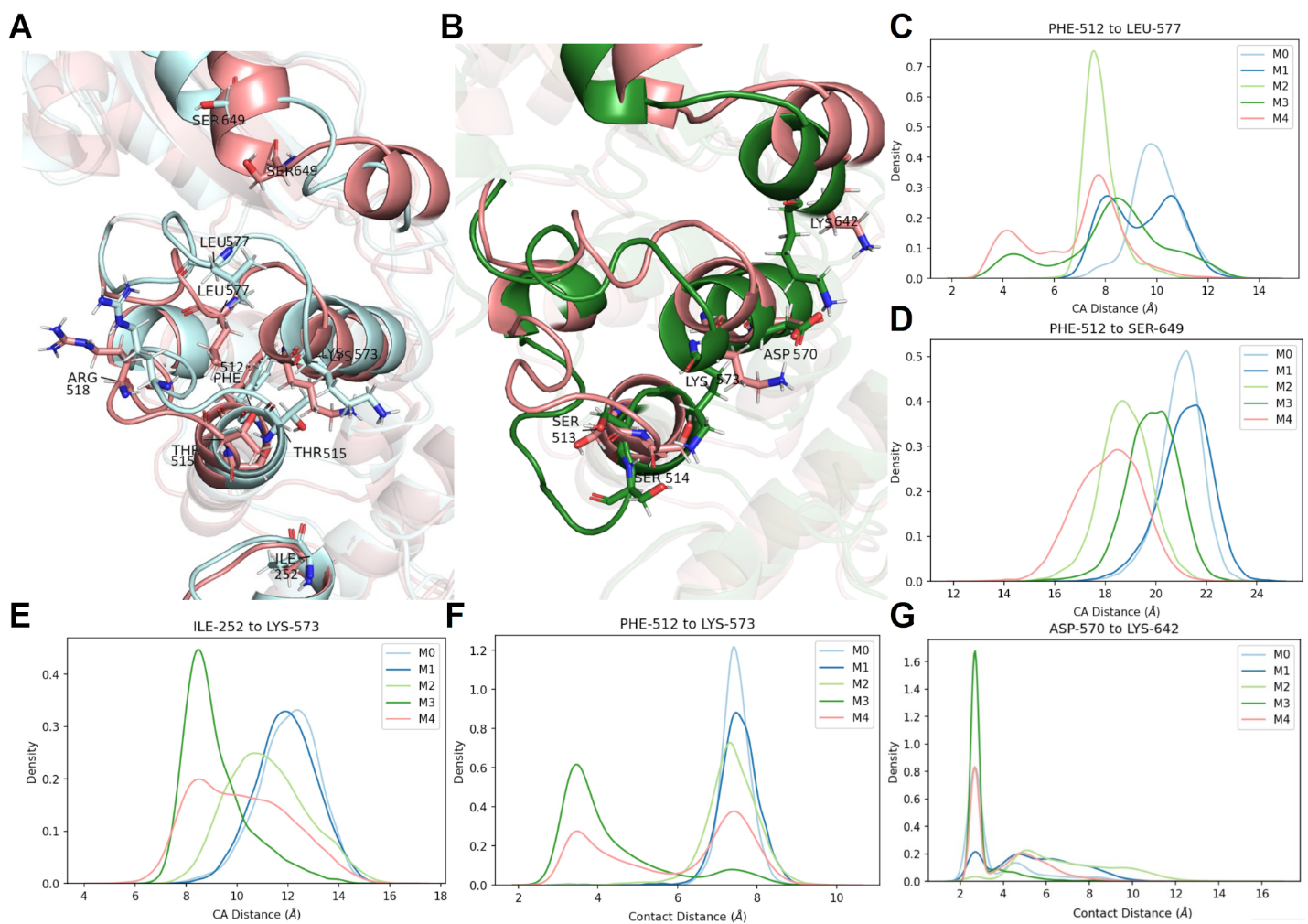

Figure S5: Proposed specific mechanism. **(A)** Representative structure macrostate 0 (light cyan) compare with macrostate 4 (salmon); **(B)** Representative structure macrostate 3 (green) compare with macrostate 4 (salmon) **(C-G)** key residue pair CA distance or contact distance (closest heavy atom distance). ILE-252 to LYS-573 distance distribution and PHE-512 to LYS-573 distace distribution show α26 shift away β22 in macrostate 0 and 1, while toward in macrostate 3; PHE-512 to LEU577 and PHE-512 to SER-649 distance distribution show α26/α27 and α32/α33 loops move away β22 in macrostate 0, 1 and 3, while in macrostate 3, α32 moves close to α26 as indicated by ASP-570 to LYS-642 distance distribution.
